# Rapid inactivation of SARS-CoV-2 variants by continuous and intermittent irradiation with a deep-ultraviolet light-emitting diode (DUV-LED) device

**DOI:** 10.1101/2021.05.10.443422

**Authors:** Hiroko Inagaki, Akatsuki Saito, Chiho Kaneko, Hironobu Sugiyama, Tamaki Okabayashi, Shouichi Fujimoto

**Affiliations:** M&N Collaboration Research Laboratory, Department of Medical Environment Innovation, Faculty of Medicine, University of Miyazaki, Japan; Department of Veterinary Science, Faculty of Agriculture, University of Miyazaki, Miyazaki, Japan; Graduate School of Medicine and Veterinary Medicine, University of Miyazaki, Miyazaki, Japan; Center for Animal Disease Control, University of Miyazaki, Miyazaki, Japan; Nikkiso Co., Ltd., Tokyo, Japan; Department of Hemovascular Medicine and Artificial Organs, Faculty of Medicine, University of Miyazaki, Japan

**Author notes:** **Corresponding author:** Shouichi Fujimoto, Department of Hemovascular Medicine and Artificial Organs, Faculty of Medicine, University of Miyazaki, Japan, 5200 Kihara, Kiyotake, Miyazaki 889-1692, Japan.

**Keywords:** SARS-CoV-2, Variants, UV-LED, Viral inactivation, COVID-19

## Abstract

More than 1 year has passed since social activities have been restricted due to the spread of severe acute respiratory syndrome coronavirus 2 (SARS-CoV-2). More recently, novel SARS-CoV-2 variants have been spreading around the world, and there is growing concern of higher transmissibility of the variants and weaker protective efficacy of vaccine against the variants. Immediate measures are needed to reduce human exposure to the virus. In this study, the antiviral efficacy of deep-ultraviolet light-emitting diode (DUV-LED) irradiation (280 ± 5 nm, 3.75 mW/cm^2^) against three SARS-CoV-2 variants was evaluated. For the B.1.1.7, B.1.351, and P.1 strains, the infectious titer reduction rates of 96.3%, 94.6%, and 91.9%, respectively, were already recognized with the irradiation of virus stocks for 1 s, and the rates increased to 99.9%, 99.9%, and 99.8%, respectively, with irradiation for 5 s. We also tested the effect of pulsed DUV-LED irradiation (7.5 mW/cm^2^, duty rate: 50%, frequency: 1 KHz) under the same output conditions as continuous irradiation, and found that the antiviral efficacy of pulsed and continuous irradiation was the same. These findings suggest that SARS-CoV-2 may be instantly inactivated by DUV-LED irradiation if the DUV-LED device is further developed and optimized to increase its output.

## Introduction

The global severe acute respiratory syndrome coronavirus 2 (SARS-CoV-2) pandemic has placed countries in a difficult and ever-evolving situation for over a year. More than 140 million cases of coronavirus disease (COVID-19) and 3.0 million deaths due to COVID-19 have been reported to the World Health Organization (WHO) as of April 18, 2021 [1]; these numbers represent increases of more than approximately 30- and 10-fold, respectively, when compared to the numbers last year. Although vaccinations have begun around the world, COVID-19 has not yet been completely suppressed. At the end of last year, three variants of SARS-CoV-2, that is, the United Kingdom (UK) strain (B.1.1.7) [2,3], South African strain (B.1.351) [4,5], and Brazilian strain (P.1) [6,7], were confirmed, and they have recently spread all over the world. These variants threaten society as a whole, since the variants may have higher transmissibility [2,3,8–10], protective efficacy of vaccine on the variants be weak [10–15] and the patients infected with the variants may be more likely to develop severe medical conditions [16,17].

However, governments worldwide are attempting to balance economic activity and medical care as much as possible. Although the development of therapeutic agents and vaccines is an important strategy for bringing an end to the pandemic, it is also necessary to devise measures to reduce virus exposure to prevent the spread of infection due to droplets and droplet nuclei.

A deep-ultraviolet light-emitting diode (DUV-LED) instrument that generates around 250- to 300-nm wavelengths has been reported to effectively inactivate microorganisms, including SARS-CoV-2 [18–22]; however, its effect on SARS-CoV-2 variants has not yet been evaluated. Recently, the inactivating effects of pulse patterns of a UV-LED device, which enables pulsed irradiation as the radiation can be turned on and off at a high frequency, on microorganisms have been shown to be as effective as continuous irradiation [23]. In this study, we examined whether continuous and intermittent (pulsed) DUV-LED irradiation can inactivate the three types of SARS-CoV-2 variants (B.1.1.7, B.1.351, and P.1).

## Materials and Methods

### Materials

#### 1. Cells

VeroE6/TMPRSS2 cells were obtained from the Japanese Collection of Research Bioresources (JCRB) Cell Bank in Japan (https://cellbank.nibiohn.go.jp/english/; JCRB no. JCRB1819). The cells were cultured in Dulbecco's Modified Eagle Medium (DMEM) containing 10% fetal bovine serum (FBS), penicillin/streptomycin, and 1 mg/ml G418 (Thermo Fisher Scientific, Tokyo).

#### 2. Viral stocks

Three types of SARS-CoV-2 variants, that is, the variants that were first described in the UK (hCoV-19/Japan/QHN001/2020 (B.1.1.7)), South Africa (hCoV-19/Japan/TY8-612/2021 (B.1.351),) and Brazil (hCoV-19/Japan/TY7-501/2020 (P.1)), were obtained from the National Institute of Infectious Diseases of Japan. These viruses were propagated in VeroE6/TMPRSS2 cells cultured in DMEM containing 10% FBS and penicillin/streptomycin. At 48 h or 72 h after infection, virus stocks were collected by centrifuging the culture supernatants at 3,000 rpm for 10 min. Clarified supernatants were kept at −80°C until use.

#### 3. DUV-LED

The DUV-LED apparatus, which generates a narrow-range wavelength (280 ± 5 nm), was obtained from Nikkiso Co. Ltd. (Tokyo, Japan). This wavelength was selected with consideration for practicality due to the high output (radiation) power and longer durability of the LED device during the developmental stage. In addition to conventional continuous irradiation, this DUV-LED instrument enables pulsed irradiation as the radiation can be turned on and off at a high frequency. We used a signal generator (AIMEX Corporation, Tokyo, Japan) to irradiate DUV-LED light.

For the evaluation of DUV-LED inactivation of the target virus, aliquots of virus stock (150 μl) adjusted to 5.0 × 10^4^ PFU/ml were placed in the center of a 60-mm Petri dish and irradiated with 3.75 mW/cm^2^ of continuous irradiation or with 7.5 mW/cm^2^ of pulsed irradiation (duty rate: 50%; frequency: 1 KHz) at a work distance of 20 mm for various times (1, 5, or 10 s; n = 3 each). We set the duty rate and frequency of the pulsed irradiation to be the same as the output per time of the continuous irradiation (Supplement 1).

### Method

The antiviral efficacy of DUV-LED irradiation against the SARS-CoV-2 variants was evaluated. After DUV-LED irradiation, approximately 120 μl of each virus stock was collected with a 200-μl tip. Virus solutions were serially diluted in 10-fold steps in serum-free DMEM in a 1.5-ml tube, then inoculated onto VeroE6/TMPRSS2 monolayers in a 12-well plate. After adsorption of virus for 2 h, cells were overlaid with MEM containing 1% carboxymethyl cellulose and 2% FBS (final concentration). The cells were incubated for 72 h in a CO_2_ incubator, then observed under a microscope for cytopathic effects. A non-irradiated virus suspension was used as a negative control. To calculate the PFU, cells were fixed with 10% formalin for 30 min, and stained with a 2% crystal violet solution.

The antiviral effects of DUV-LED irradiation were assessed using the logPFU ratio calculated as log10 (Nt / N0), where Nt is the PFU count of the UV-irradiated sample, and N0 is the PFU count of the sample without UV irradiation. In addition, the infectious titer reduction rate was calculated as (1 − 1 / 10log PFU ratio) × 100 (%). All experiments were performed in a biosafety level 3 laboratory.

## Results

### Inactivating effects of continuous irradiation with a DUV-LED device

We observed a marked cytopathic effect in all cells that were infected with the UK, South African, or Brazilian strain and were not irradiated with DUV-LED light (Figure 1a). The infected cells that were irradiated for 1 s showed an obvious reduction in the cytopathic effect (Figure 1b), and the morphology of the cells that were irradiated for 5 s was largely comparable to that of the mock cells (Figure 1c, d).

**Figure 1.**
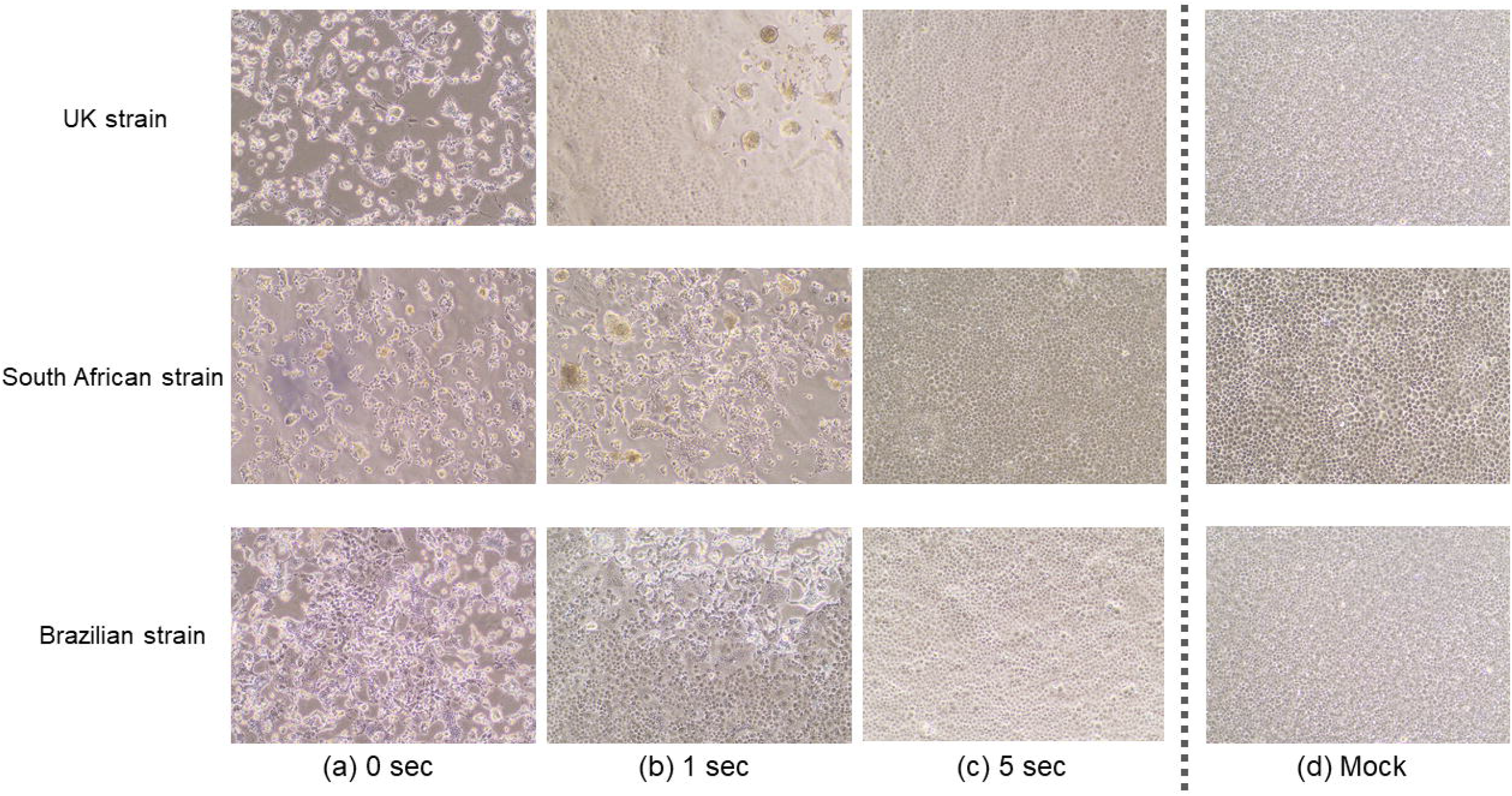
Inhibitory effects of continuous DUV-LED irradiation on three types of SARS-CoV-2 variants (the UK, South African, and Brazilian strains). Cytopathic changes in virus-infected VeroE6/TMPRSS2 cells without DUV-LED irradiation (a), with DUV-LED irradiation for 1 s (b) or 5 s (c) corresponding to 3.75 or 18.75 mJ/cm^2^, respectively. (d) Mock cells.

The plaque assay revealed that a short irradiation time inactivated SARS-CoV-2 variants rapidly (Figure 2). Of note, for the UK, South African, and Brazilian strains, the infectious titer reduction rates of 96.3%, 94.6%, and 91.9%, respectively, were already recognized with the irradiation of virus stocks for 1 s, and the rates increased to 99.9%, 99.9%, and 99.8%, respectively, with irradiation for 5 s (Table 1 and Figure 3). These results suggested that DUV-LED irradiation for a very short time can drastically inactivate SARS-CoV-2 variants.

**Figure 2.**
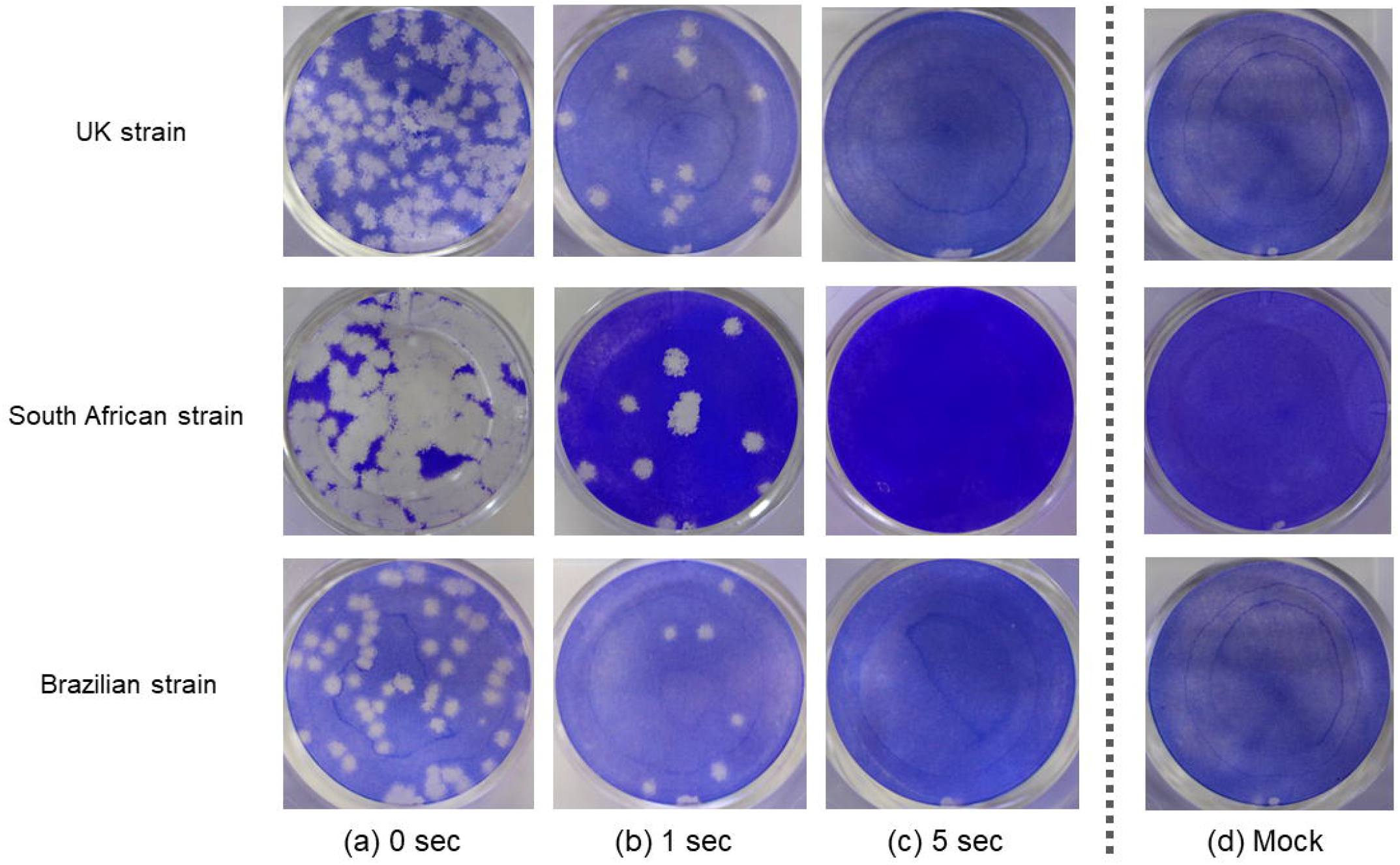
Plaque formation in VeroE6/TMPRSS2 cells. Virus solutions were treated with continuous DUV-LED irradiation for 0, 1, or 5 s, then diluted 100-fold and inoculated onto VeroE6/TMPRSS2 cells. Representative results are shown.

**Table 1.** Differences in the infectious titer after continuous and pulsed DUV-LED irradiation for the UK, South African, and Brazilian strains irradiated with different patterns of DUV-LED light for 0, 1, 5, or 10 s.

|  |  | Control<br>(no irradiation) | DUV-LED irradiation time |  |  |  |  |  |
| --- | --- | --- | --- | --- | --- | --- | --- | --- |
|  |  |  | 1 sec |  | 5 sec |  | 10 sec |  |
|  |  |  | continuous<br>irradiation | pulsed<br>irradiation | continuous<br>irradiation | pulsed<br>irradiation | continuous<br>irradiation | pulsed<br>irradiation |
| UK strain | PFU (PFU/mL) | $3.5 \times 10^4$ | $1.3 \times 10^3$ | $1.9 \times 10^3$ | < 20 | < 20 | < 20 | < 20 |
|  | Log PFU ratio<sup>a</sup> | — | 1.4 | 1.3 | > 3.2 | > 3.2 | > 3.2 | > 3.2 |
|  | Infection titer reduction ratio<sup>b</sup> (%) | — | 96.3 | 94.4 | > 99.9 | > 99.9 | > 99.9 | > 99.9 |
| South African strain | PFU (PFU/mL) | $5.3 \times 10^4$ | $2.9 \times 10^3$ | $3.5 \times 10^3$ | < 20 | $5.3 \times 10^1$ | < 20 | < 20 |
|  | Log PFU ratio<sup>a</sup> | — | 1.3 | 1.2 | > 3.4 | 3.1 | > 3.4 | > 3.4 |
|  | Infection titer reduction ratio<sup>b</sup> (%) | — | 94.6 | 93.4 | > 99.9 | 99.9 | > 99.9 | > 99.9 |
| Brazilian strain | PFU (PFU/mL) | $1.1 \times 10^4$ | $8.7 \times 10^2$ | $1.7 \times 10^3$ | < 20 | < 20 | < 20 | < 20 |
|  | Log PFU ratio<sup>a</sup> | — | 1.1 | 0.8 | > 2.7 | > 2.7 | > 2.7 | > 2.7 |
|  | Infection titer reduction ratio<sup>b</sup> (%) | — | 91.9 | 84.4 | > 99.8 | > 99.8 | > 99.8 | > 99.8 |

<sup>a</sup> $\log_{10} (N_t/N_0)$ where $N_t$ is the PFU count of the UV-irradiated sample and $N_0$ is the PFU count of the sample without UV irradiation.
<sup>b</sup> $(1 - 1/\log_{10} \text{PFU ratio}) \times 100$ (%).

**Figure 3.**
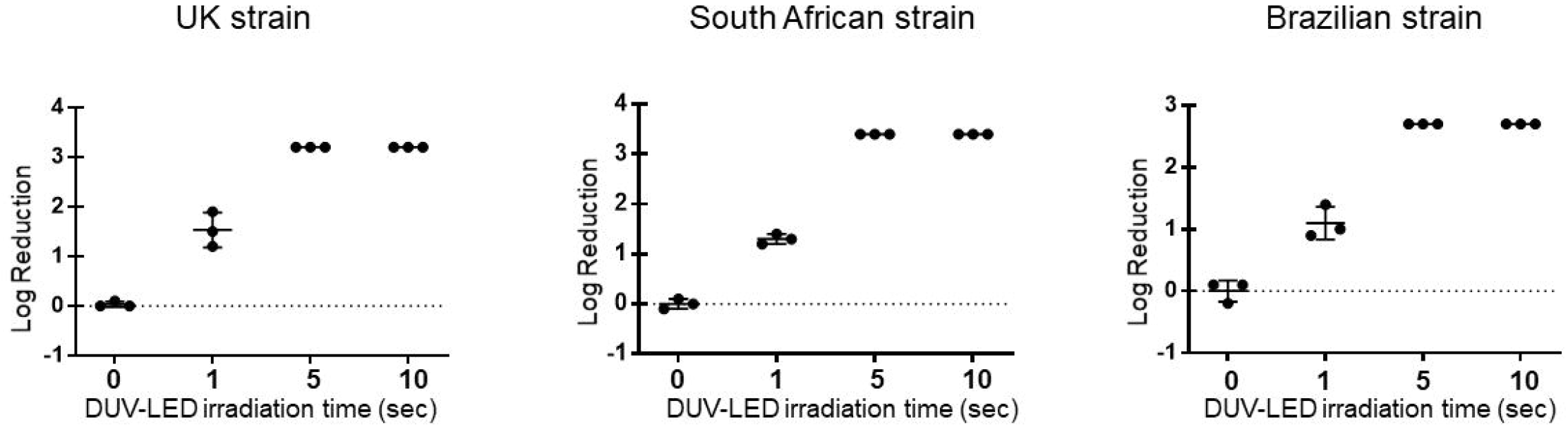
Log reduction of the infectious titer by continuous irradiation with DUV-LED for the UK, South African, and Brazilian strains. DUV-LED: continuous irradiation (current: 0.35 A). Time-dependent inactivation of SARS-CoV-2 by DUV-LED irradiation. The results shown are the means and standard deviations of triplicate measurements.

### Inactivating effects of pulsed irradiation with a DUV-LED device

For the UK, South African, and Brazilian strains, the infectious titer reduction rates of 94.4%, 93.4%, and 84.4%, respectively, were already recognized with the irradiation of virus stocks for 1 s, and the rates increased to 99.9%, 99.9%, and 99.8%, respectively, with irradiation for 5 s (Table 1 and Figure 4). These results were almost the same as those obtained with continuous irradiation.

**Figure 4.**
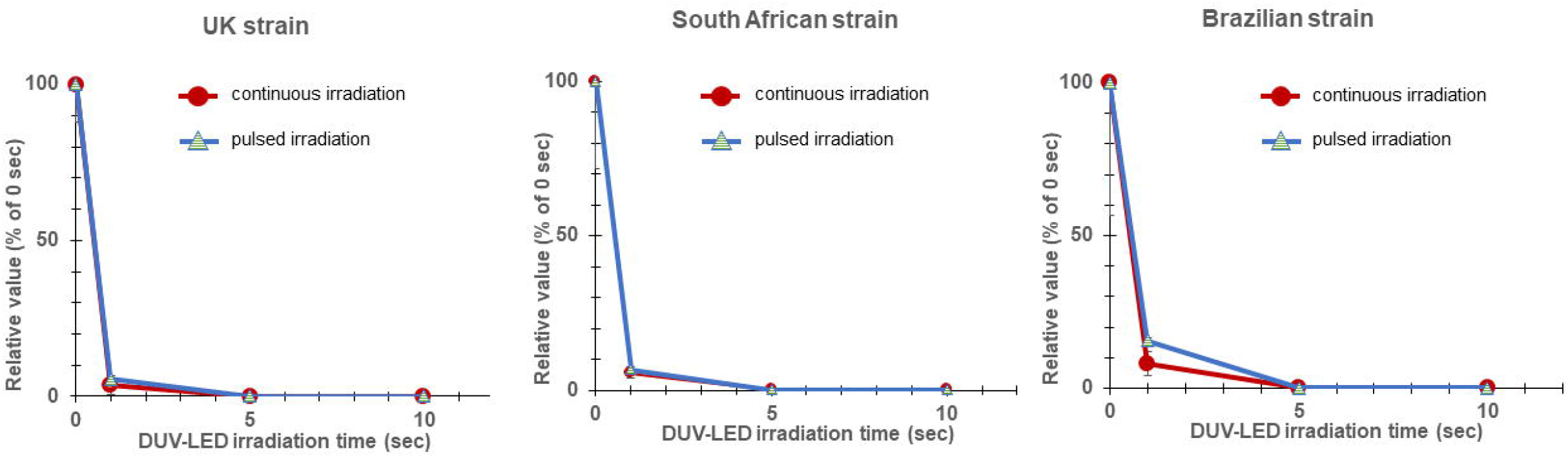
Relative value (percentage of the value of the cells irradiated for 0 s) of the infectious titer after continuous and pulsed irradiation with DUV-LED for the UK, South African, and Brazilian strains. DUV-LED: continuous irradiation (current: 0.35 A) or pulsed irradiation (current: 0.7 A; duty ratio: 50%; frequency: 1 KHz). The inactivating effects were almost the same between the continuous and pulsed DUV-LED irradiation.

## Discussion

Recently, there has been growing concern around the world, including at the WHO, regarding the emergence of new SARS-CoV-2 variants since they may have longer infectivity periods [24], stronger infectivity, including in children [2,8,10,25,26], and protective efficacy of vaccine on these variants may be weak [12–15]. It has also been pointed out that patients infected with SARS-CoV-2 variants may be more likely to develop severe medical conditions and have a higher risk of death [8,10,16]. Therefore, in parallel with advances in vaccines and therapeutic agents, measures to reduce virus exposure to prevent the spread of SARS-CoV-2 infection are desired.

The present study demonstrated for the first time that DUV-LED irradiation can rapidly inactivate three types of SARS-CoV-2 variants, that is, the ones that were first described in the UK, South Africa, and Brazil [2–7,9,10,26]. Additionally, continuous and pulsed DUV-LED irradiation showed similar degrees of rapid virus inactivation.

UV-LED devices that can provide irradiation at various peak emission wavelengths, such as UV-A (320 − 400 nm), UV-B (280 − 320 nm), and UV-C (100 − 280 nm), have been adopted to inactivate various pathogenic species, including bacteria, viruses, and fungi. UV-C is considered to be the most effective germicidal region of the UV spectrum as it causes the formation of photoproducts in DNA and RNA [27]. These pyrimidine dimers interrupt the transcription, translation, and replication of DNA and RNA, and eventually lead to the death of the microorganism [28]. Last year, we reported for the first time that irradiation with DUV-LED at a wavelength of 280 ± 5 nm rapidly inactivated wild-type SARS-CoV-2 that was obtained from a COVID-19 patient [22]. The effect of DUV-LED irradiation on the wild-type SARS-CoV-2 (infectious titer reduction rates of 87.4% for 1 s and 99.9% for 10 s of irradiation) was similar to that on the SARS-CoV-2 variants in the present study. Since the SARS-CoV-2 variants also have a lipophilic outer membrane (envelope protein), the inactivating effects of DUV-LED irradiation on the UK, South African, and Brazilian variants were as expected, and similar effects are also expected for other variants that may emerge in the future. Unlike enveloped viruses, non-enveloped viruses that do not have the envelope protein, such as norovirus, are known to be highly resistant to disinfectants, but DUV-LED irradiation is expected to be effective against non-enveloped viruses as well (unpublished observations).

In this study, we also tested the effect of pulsed irradiation with a DUV-LED device. As shown in Table 1 and Figure 4, the degree of virus inactivation by continuous and intermittent irradiation was comparable when the device outputs were the same (power X radiation time). This suggested that SARS-CoV-2 may be instantly inactivated by DUV-LED irradiation if the DUV-LED device is further developed and optimized to increase its output [23].

Despite the significant inactivating effects of DUV-LED reported here, this study has some limitations. First, these effects may be limited to the test conditions applied in this study, including the irradiation distance and output (work distance of 20 mm; irradiation at 3.75 mW/cm^2^ for continuous irradiation and 7.5 mW/cm^2^ for pulsed irradiation (duty rate: 50%; frequency: 1 KHz)). In addition, it is necessary to also evaluate multiple parameters, such as the frequency and duty ratio, to clarify the effectiveness of pulse irradiation. The influence of the material and the power consumption for the high amplitude were also not evaluated. In the future, we will examine in more detail whether various conditions of DUV-LED irradiation may affect the degree of the inactivation of microorganisms.

In addition to community settings, healthcare settings are also vulnerable to the invasion and spread of SARS-CoV-2 and its variants, and the stability of SARS-CoV-2 in aerosols and on surfaces [19] likely contributes to the transmission of the virus in medical environments. It is important to create an environment that minimizes virus exposure to suppress the spread of SARS-CoV-2 in a sustainable and efficient manner. It was confirmed in our study that SARS-CoV-2, including its variants, is highly susceptible to DUV-LED irradiation. By devising appropriate and optimized irradiation methods, it is conceivable that DUV-LED irradiation can be adapted and applied in various settings. This study provides useful baseline data for securing a safer community and medical environment. The development of devices equipped with DUV-LED is expected to prevent virus spread through the air and from contaminated surfaces.

## Contributors

H.I., H.S. and A.S. conceived the study and wrote the manuscript. A.S., C.K., and T.O. conducted the experiments dealing with the viruses. S.F. contributed to the study design, study supervision, and manuscript revision.

## Acknowledgements

We wish to thank the National Institute of Infectious Diseases, Japan, for providing the hCoV-19/Japan/TY7-501/2020, hCoV-19/Japan/TY8-612/2021, and hCoV-19/Japan/QHN001/2020 strains. This study was supported in part by the Japan Agency for Medical Research and Development Research Program on Emerging and Re-emerging Infectious Diseases (20fk0108163 and 20fk0108518 to A.S.); Japan Agency for Medical Research and Development Japan Program for Infectious Diseases Research and Infrastructure (20wm0325009 to A.S.); Japan Society for the Promotion of Science (JSPS) KAKENHI Grant-in-Aid for Scientific Research (B) (21H02361 to T.O. and A.S.); JSPS KAKENHI Grant-in-Aid for Scientific Research (C) (19K06382 to A.S.); and JSPS KAKENHI Grant-in-Aid for Early-Career Scientists (19K15984 to C.K.); the Grant for Joint Research Project of the Research Institute for Microbial Diseases, Osaka University (to T.O. and A.S.).

## Supplement caption

**Supplement 1.**
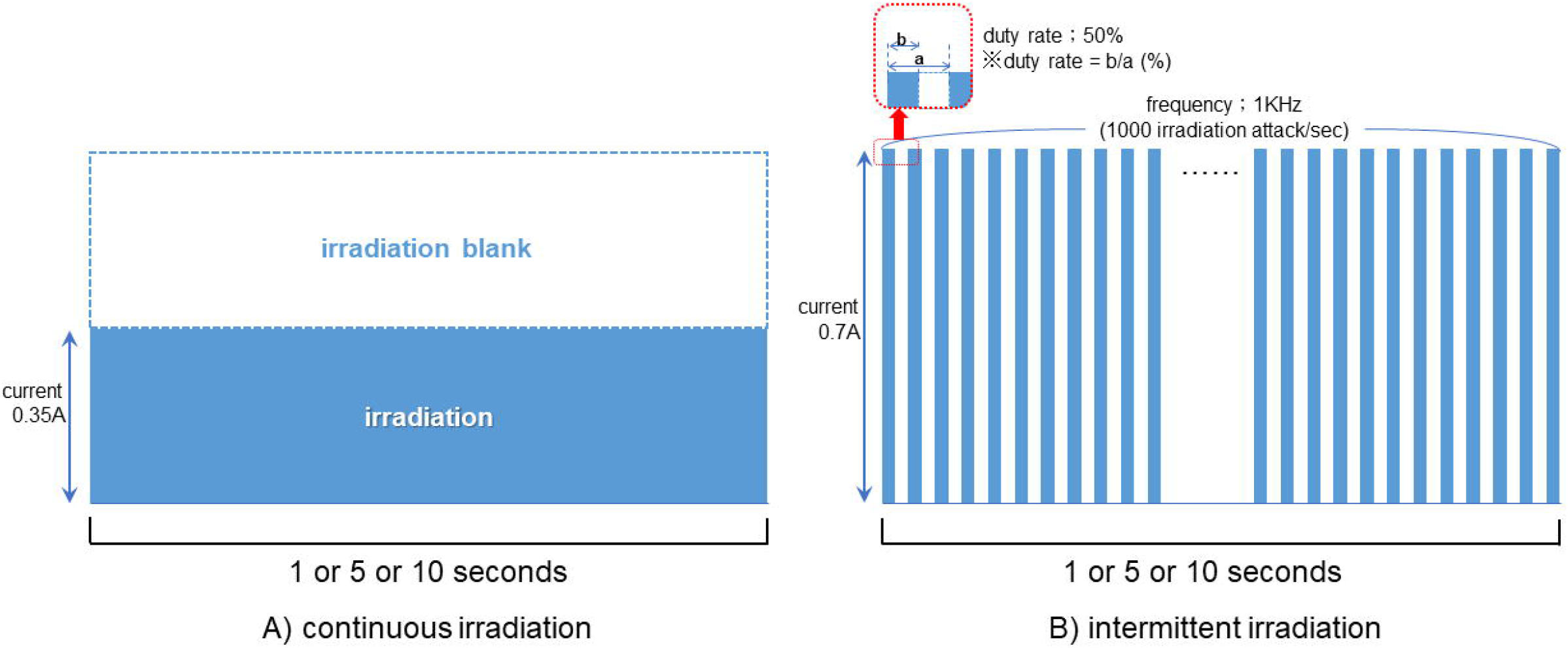
Output diagrams of two patterns of DUV-LED irradiation. A) Continuous irradiation with a current of 0.35 A for 1, 5, or 10 s. B) Intermittent (pulsed) irradiation with a current of 0.7 A for 1, 5, or 10 s (frequency: 1 KHz; duty rate: 50%). The output per time was set to be the same in continuous irradiation (A, 3.75 mW/cm^2^ × time) and intermittent irradiation (B, 7.5 mW/cm^2^ × time).

